# Very small spontaneous structural mutation rate in green algae

**DOI:** 10.1101/2022.02.09.479524

**Authors:** Matéo Léger-Pigout, Marc Krasovec

## Abstract

Structural mutations led to major innovations, but our knowledge of the spontaneous structural mutation rate is very limited. We cannot have a complete view of the adaptive potential of species and new variants in a population without addressing this question. Here, we used Illumina genomic data from mutation accumulation experiments of five green algae species and measured very low structural mutation rate, between 5×10^-12^ to 2×10^-11^ mutations per site, at least ten times lower than the nucleotide mutation rate in these species. Such low rates of structural mutations may be a consequence of selection. Assuming that structural mutations have higher fitness effect than point mutations because they impact a larger portion of the genome, selection toward a small structural mutation rate may be stronger than for point mutations. A second explanation could be the rate of true positive detection of de novo structural mutations, that is globally poor and variable between methods. This highlights the necessity of long read sequencing in future mutation accumulation studies.

**SIGNIFICANCE STATEMENT:** Mutations are the first source of all biodiversity. Biologists tried since a century to measure the spontaneous mutation rate, but our knowledge on the structural spontaneous mutation rate, *μ_st_*, is still poor. Here we measured *μ_st_* in five species of phytoplankton, and found a very small rate compared to nucleotide and short insertion-deletion mutations.

## INTRODUCTION

Spontaneous mutations renew the standing genetic variation and permit adaptation through natural selection. The spontaneous mutation rate *μ* is so one of the most important parameters to understand evolution. Since decades, biologists tried to measure it using mutation accumulation (MA) experiment (Halligan & Keightley, 2009), first with a phenotypic approach by following fitness variation of MA lines (Mukai, 1964), and then directly by whole genome sequencing (Lynch *et al.*, 2016; Katju & Bergthorsson, 2019). Our knowledge made significant progresses the last fifteen years, particularly on the nucleotide (Lynch *et al.*, 2016; Katju & Bergthorsson, 2019) and short insertion-deletion (Sung *et al.*, 2016) mutation rates, noted *μ_bs_* and *μ_id_*, respectively. We observe 3-fold variation between species, from ~1×10^-11^ to ~1×10^-8^ mutations per site per haploid genome in ciliates and humans. This variation is in part explained by the drift barrier hypothesis (Sung *et al.*, 2012) where large effective population size (*N_e_*) species reaches lower *μ* than small *N_e_* species. As a consequence of the mutation load, selection pushes toward a small *μ*, that is more efficient in large *N_e_* where drift is limited. This hypothesis is currently the most accepted, even if other factors play on the mutation rate but with smaller effect such as GC content (Krasovec *et al.*, 2017), genome size, generation time or metabolic rate (Martin & Palumbi, 1993; Mooers & Harvey, 1994; Thomas *et al.*, 2010; Weller & Wu, 2015). Both *μ_bs_* and *μid* depends so mostly of *N_e_*, but our knowledge about the spontaneous structural mutation rate *μ_st_* is very poor (Press *et al.*, 2019; Ho & Schaack, 2021) and available in only few models (see Table 3 of Katju and Bergthorsson 2019 for copy number variant for example). Structural mutations, defined here as duplications, large insertions-deletions, inversions or chromosome rearrangements, may have stronger phenotypic effect than point mutations because they impact a larger portion of the genome. Thus, the pressure toward a small *μ_st_* should be stronger than for *μ_bs_* and *μ_id_*. In addition, it is impossible to have a complete view of new variants and adaptive potential of a species with limited knowledge of *μ_st_*. Structural mutations led to major innovations, for example the supergenes following inversions. In birds, this process saw the emergence of different morphotypes in a same species with different colors, behaviors and feather shapes (Küpper *et al.*, 2016; Tuttle *et al.*, 2016). Here, we measured *μ_st_* for five species of green algae using mutation accumulation experiment Illumina reads from previous publications (Krasovec *et al.*, 2016, 2017, 2018).

## MATERIALS AND METHODES

Illumina raw reads of 150 and 300 bp paired-end were mapped to the reference genomes with bwa mem (Li & Durbin, 2010), then bam files were treated with samtools (Li *et al.*, 2009) and structural variant called with delly (Rausch *et al.*, 2012), svaba (Wala *et al.*, 2018) and lumpy (Layer *et al.*, 2014) with standard parameters (excepted ploidy at 1). For svaba, we used -n option to input the T_0_ genome bam file as control. VCF files were then sorted with several thresholds to increase the rate of true de novo mutations identification: the mutated site must be covered by 10 in the concerned MA line and the T0 genome; the mapping quality of the site must be higher than 20; the variant must be present in only one MA line, expected for few variants present in two or three MA lines because of non-independent lines (in that case common mutations and generations were count as one); the variant must totalize all the coverage of the site; last, some structural variants appeared only in one MA line, but were not count as mutation because other lines had same variants but at different positions within the confidence interval of start-end positions. De novo structural mutations candidates were then checked by coverage with samtools and by PCR and Sanger sequencing when possible.

## RESULTS AND DISCUSSION

### False - true positive rate of structural de novo mutation detection

Twenty-six de novo structural mutation candidates were found in the 5 species with 3 softwares (Table 1), but coverage and PCR checks reduced this number to 1, given a very low true positive rate of 4%. From Sanger sequencing, the 8 tested breaks from svaba and delly were false, so we considered all other non-tested breaks false. The other detected candidates were deletions and duplications, all larger than 300 pb apart one deletion of 215 pb. According to Wala et al (Wala *et al.*, 2018), the positive rates of deletions larger than 300 bp are 85% for svaba and delly, and 95% for lumpy; for insertion/duplication, it decreases to 37%, 24% and 29%. Here, the positive rate of deletion is 20% and the 3 duplication candidates were false. The positive rate of duplications was so 0%, but with only 3 candidates this rate should be taken with caution. Such low positive rates show the difficulties to identify structural de novo mutations with good accuracy, particularly with short reads. The de novo mutation candidates were different between the three softwares, and the single true positive was found with lumpy only, which implies that some true mutations have been possibly missed. Further investigations of mutation accumulation experiments coupled with long reads sequencing should be the next milestone to significantly increase the detection accuracy.

**Table 1.** List of de novo structural mutation candidates. Only the deletion of Om21b is true.

| MA line | Mutation type | Software | Start | End | PCR - Sanger | Coverage |
| --- | --- | --- | --- | --- | --- | --- |
| Ot20 | break | delly | NC_014444.2: 184889 | NC_014431.2: 783883 | false | na |
| Om14 | break | delly | BCC102-800924: 233 | BCC102-1186077: 1181287 | false | na |
| Om17 - 18 | break | delly | BCC102-594986: 60163 | BCC102-768578: 224081 | false | na |
| Om1 | break | delly | BCC102_ML_SOC: 68272 | BCC102-397956: 160264 | false | na |
| Om2 | break | delly | BCC102-466838: 2048 | BCC102-867403: 392 | false | false |
| Ot38 | deletion | delly | NC_014427.2: 430027 | NC_014427.2: 438281 | no check | false |
| Bp8 | deletion | delly | NC_023992.1: 297114 | NC_023992.1: 301342 | false | false |
| Bp20 | duplication | delly | NC_023992.1: 296522 | NC_023992.1: 297150 | false | false |
| Ot38 | break | svaba | NC_014427.2: 820183 | NC_014428.2: 994946 | false | na |
| Ot38 | break | svaba | NC_014433.2: 406614 | NC_014437.2: 310849 | false | na |
| Om13 | break | svaba | BCC102-873724: 213431 | BCC102_ML_SOC: 100063 | false | na |
| Ot22 | break | lumpy | NC_014426.2: 181294 | NC_014428: 229429 | no check | na |
| Ot23 | break | lumpy | NC_014426.2: 929056 | NC_014442.2: 399570 | no check | na |
| Ot9 - 10 | break | lumpy | NC_014435.2: 273589 | NC_014439.2: 385664 | no check | na |
| Ot20 - 31 | break | lumpy | NC_014441.2: 17675 | NC_014443.2: 58813 | no check | na |
| Om38 | break | lumpy | BCC102-993683: 535232 | BCC102-768578: 224085 | no check | na |
| Om12 | break | lumpy | BCC102-637367: 375185 | BCC102-637367: 319399 | no check | na |
| Om3 | break | lumpy | BCC102-873724: 329213 | BCC102-768578: 224085 | no check | na |
| Om13 | break | lumpy | BCC102-873724: 173257 | BCC102-546026: 350305 | no check | na |
| Om28 | break | lumpy | BCC102-867403: 378390 | BCC102-496340: 275198 | no check | na |
| Mp27 - 34 - 35 | break | lumpy | chrom_01: 564769 | chrom_01: 1360407 | no check | na |
| Om9 - 29d | deletion | lumpy | BCC102-867403: 33955 | BCC102-867403: 34170 | false | false |
| Om31 | deletion | lumpy | BCC102-466838: 274207 | BCC102-466838: 276502 | false | false |
| Om21b | deletion | lumpy | BCC102_ML_SOC: 571078 | BCC102_ML_SOC: 572460 | true | true |
| Om17 | duplication | lumpy | BCC102-806365: 187888 | BCC102-806365: 188243 | no check | false |
| Om32 | duplication | lumpy | BCC102_ML_SOC: 243207 | BCC102_ML_SOC: 251990 | no check | false |

### *μ_st_* is more than 10 time lower than *μ_bs_* and *μ_id_*

The single de novo mutation validated by PCR is a deletion of 1,382 bp identified in *Ostreoccocus mediterraneus* on the outlier chromosome (Yau *et al.*, 2020), a particular chromosome with low GC content. The mutated region is intergenic and the deleted sequence appears elsewhere in the genome (Table S1). This makes this deleted sequence a possible repeat region resulting from transposable elements, commonly known to be implicated in this type of mutation. This single mutation gives a very low *μ_st_* for the 5 species, between 5×10^-12^ and 2×10^-11^ structural mutations per site per haploid genome per generation (Figure 1), ten times lower than *μ_bs_* and *μ_id_* in these species. We see three explanations for such low structural mutation rate. First is the poor sensitivity of the method detection (see above), resulting in an artificially low structural mutation rate. Second, low structural mutation rate may come from the mutation mechanisms themself. Several mechanisms act directly on nucleotides such as methylation and oxidation and are known to increase *μ_bs_*, making it higher than *μ_st_* or *μ_id_* just because single sites have more chance to mutate. Third is selection, for which efficiency depends of *N_e_* estimated around few millions to 10 million in phytoplankton species (Blanc-Mathieu *et al.*, 2017; Krasovec *et al.*, 2019, 2020). With such small *μ_st_*, we may argue that selection pushes harder toward a low *μ_st_* than for *μ_bs_* and *μ_id_*. This is propable assuming that structural mutations have stronger and more frequent deleterious effects than point mutations because they change a larger part of the genome. The genomes of species studied here are small, very compacted with few introns and small distance between genes. For example, *O. tauri* has a genome size of 12.5 Mb, with 81.6% of coding sequences, 82% of genes without introns and an average distance between genes of 254 bp. In that case, any structural mutation has high chance to touch at least one gene or its regulatory region, making almost all structural mutations deleterious. We can hypothesis that it is less the case in larger genomes with a longer distance between genes and high proportion of repeats and non-coding sequences.

**Figure 1.**
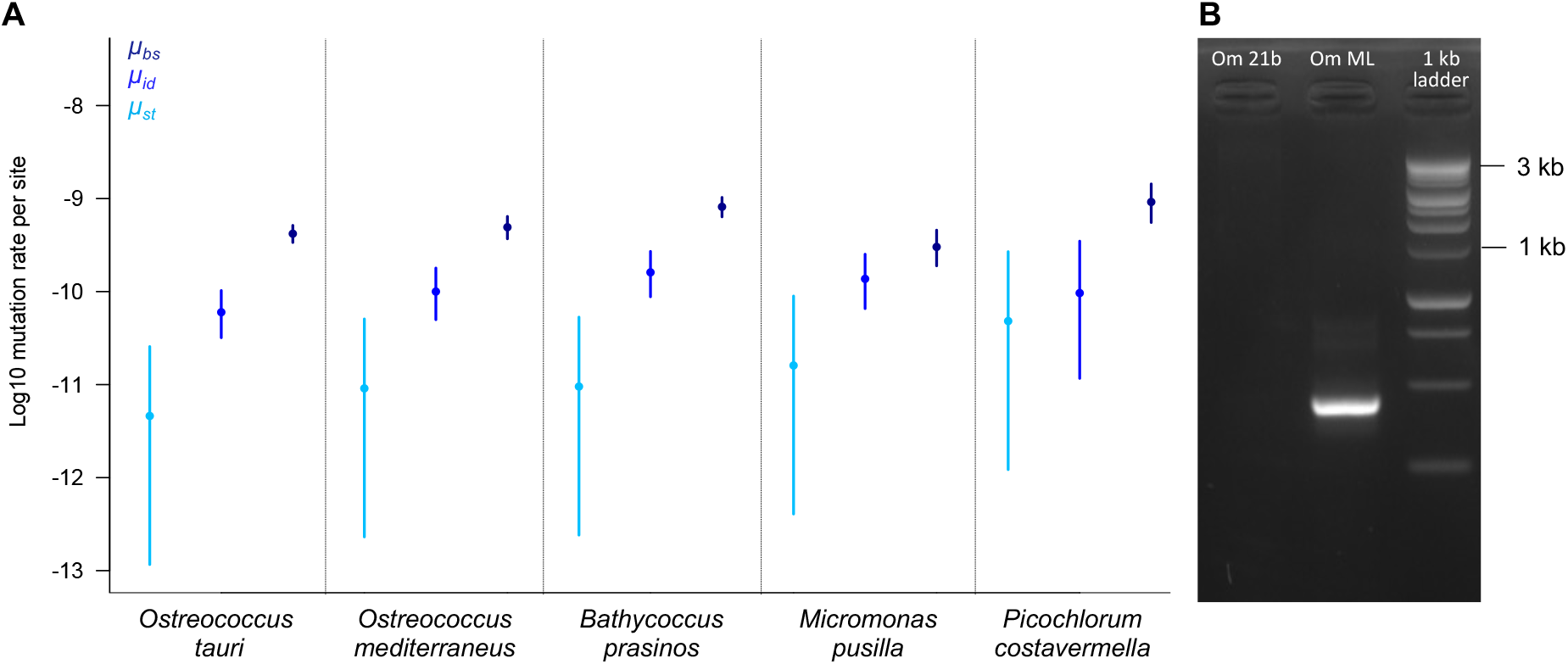
**A**. Spontaneous mutation rate per nucleotide per haploid genome per generation. Light blue: *μ_st_*, spontaneous structural mutation rate. Blue: *μ_id_*, spontaneous nucleotide mutation rate. Dark blue: *μbs*, spontaneous short insertion-deletion mutation rate. Horizontal bars are confidence interval following a Poisson distribution. *μ_id_* and *μ_bs_* come from previous studies (Krasovec *et al.*, 2017, 2018). *μ_st_* for *O. tauri*, *M. pusilla*, *B. prasinos* et *P. costavermella* are calculated assuming one hypothetical mutation (non has been validated). **B.** PCR product migration of one large deletion (1,382 bp in intergenic region) identified in *O. mediterraneus*. The ladder is the 1 kb DNA ladder from Promega. No amplification is observed in the mutation accumulation line Om 21b because primers were designed insight the deleted sequence. OmML is the T0 genome of the mutation accumulation.

## DATA AVAILABILITY STATEMENT

All genomic raw reads used in this study are available on the SRA database in the NCBI under the bioprojects PRJNA531882 (*Ostreoccocus tauri*, *O. mediterraneus*, *Micromonas pusilla*, *Bathycoccus prasinos*), PRJNA453760 and PRJNA389600 (*Picochlorum costavermella*).

## ACKNOWLEDGEMENTS

We are grateful to all GENOPHY team of the Banyuls sur mer lab for discussions and advices.

## AUTHOR CONTRIBUTIONS

MK designed and managed the study, MLP did bioinformatics analysis and wet lab works, MK wrote the draft of the manuscript, all authors contributed to editing the last manuscript version.

---

**Table S1.** Blastn results of the deleted sequence in *Ostrecoccus mediterraneus*. The blasted sequence was the sequence amplified by PCR in the MA mother line (404 bp length).

| Chromosome | Identity | Length | Start | End | Pvalue |
| --- | --- | --- | --- | --- | --- |
| BCC102-546026 | 98.916 | 369 | 358951 | 359319 | 0.0 |
| BCC102-546026 | 93.590 | 156 | 356541 | 356696 | 3.17e-62 |
| BCC102-546026 | 89.062 | 128 | 468750 | 468877 | 5.46e-40 |
| BCC102-466838 | 91.617 | 167 | 435797 | 435963 | 9.76e-62 |
| BCC102_ML_SOC | 99.010 | 404 | 321019 | 321422 | 0.0 |
| BCC102_ML_SOC | 98.721 | 391 | 571052 | 571442 | 0.0 |
| BCC102_ML_SOC | 90.678 | 354 | 225814 | 226166 | 2.59e-133 |
| BCC102_ML_SOC | 90.571 | 350 | 257243 | 257591 | 4.34e-131 |
| BCC102_ML_SOC | 90.571 | 350 | 584735 | 585083 | 4.34e-131 |
| BCC102_ML_SOC | 90.299 | 134 | 208824 | 208955 | 2.28e-44 |

